# Exogenous fatty acids inhibit fatty acid synthesis through competition between endogenously- and exogenously-generated substrates for phospholipid synthesis in *Escherichia coli*

**DOI:** 10.1101/2024.10.28.620573

**Authors:** Stefan Pieter Hendrik van den Berg, Adja Zoumaro-Djayoon, Flora Yang, Gregory Bokinsky

## Abstract

Exogenous fatty acids are directly incorporated into bacterial membranes, heavily influencing bacterial ecology and antibiotic susceptibility. We use liquid chromatography/mass spectrometry to characterize how exogenous fatty acids impact the *Escherichia coli* fatty acid synthesis pathway. We find that acyl-CoA synthesized from exogenous fatty acids rapidly increases long-chain acyl-ACP levels while depleting malonyl-ACP, indicating inhibition of fatty acid synthesis. Contrary to previous assumptions, acyl-CoA does not inhibit FabI in vivo; instead, substrate competition between acyl-CoA and acyl-ACP for phospholipid synthesis enzymes causes long-chain acyl-ACP to accumulate, inhibiting fatty acid synthesis initiation. Furthermore, changes in the acyl-ACP pool driven by acyl-CoA amplify the effects of exogenous fatty acids on the balance between saturated and unsaturated membrane lipids. Transcriptional regulation rebalances saturated and unsaturated acyl-ACP by adjusting FabA and FabB expression. Remarkably, all other fatty acid synthesis enzymes remain at stable levels, maintaining a fixed synthesis capacity despite the availability of exogenous fatty acids. Since all bacterial pathways for exogenous fatty acid incorporation characterized so far converge with endogenous synthesis pathways in a common substrate pool, we propose that the substrate competition-triggered feedback mechanism identified here is ubiquitous across bacterial species.

## Introduction

The synthesis of lipids for membrane biogenesis consumes considerable amounts of energy and metabolites. Microorganisms avoid this cost whenever possible by incorporating exogenous fatty acids directly into the cellular membrane (reviewed in (1)). Exogenous fatty acids exert profound effects on bacterial physiology and ecology and are particularly relevant to infectious disease. In pathogenic bacteria, exogenous fatty acids can facilitate or impede cold resistance (2, 3), influence biofilm formation, and increase membrane permeability and antibiotic susceptibility (4, 5). Recently, host-derived fatty acids were found to support healthy microbiota within the female genital tract and proved more effective than antibiotics *in vitro* at promoting strains that prevent bacterial vaginosis (6). Exogenous fatty acids enable some bacteria to evade fatty acid synthesis inhibitors (7), a finding that triggered an important debate about whether bacterial fatty acid synthesis is an appropriate target for clinical antibiotics. However, very few species are able to entirely replace fatty acid synthesis with exogenous fatty acids (1, 8). In many organisms, fatty acid synthesis is needed to supply precursors for essential membrane components such as lipopolysaccharides. *De novo-* synthesized lipids also maintain membrane properties such as fluidity. How organisms regulate *de novo* fatty acid synthesis in response to exogenous fatty acid incorporation is incompletely understood.

In the model species *Escherichia coli,* long-chain fatty acids are imported via the membrane channel FadL (9, 10) and activated by the long-chain fatty acid-CoA ligase enzyme FadD (11). A fraction of the acyl-CoA pool is then catabolized into acetyl-CoA by the fatty acid degradation pathway, while acyl-CoA of appropriate length are incorporated into phospholipids by the glycerol-3- phosphate acyl transferase PlsB and the lysophosphatidic acid acyl transferase PlsC (**Figure 1**).

**Figure 1.**
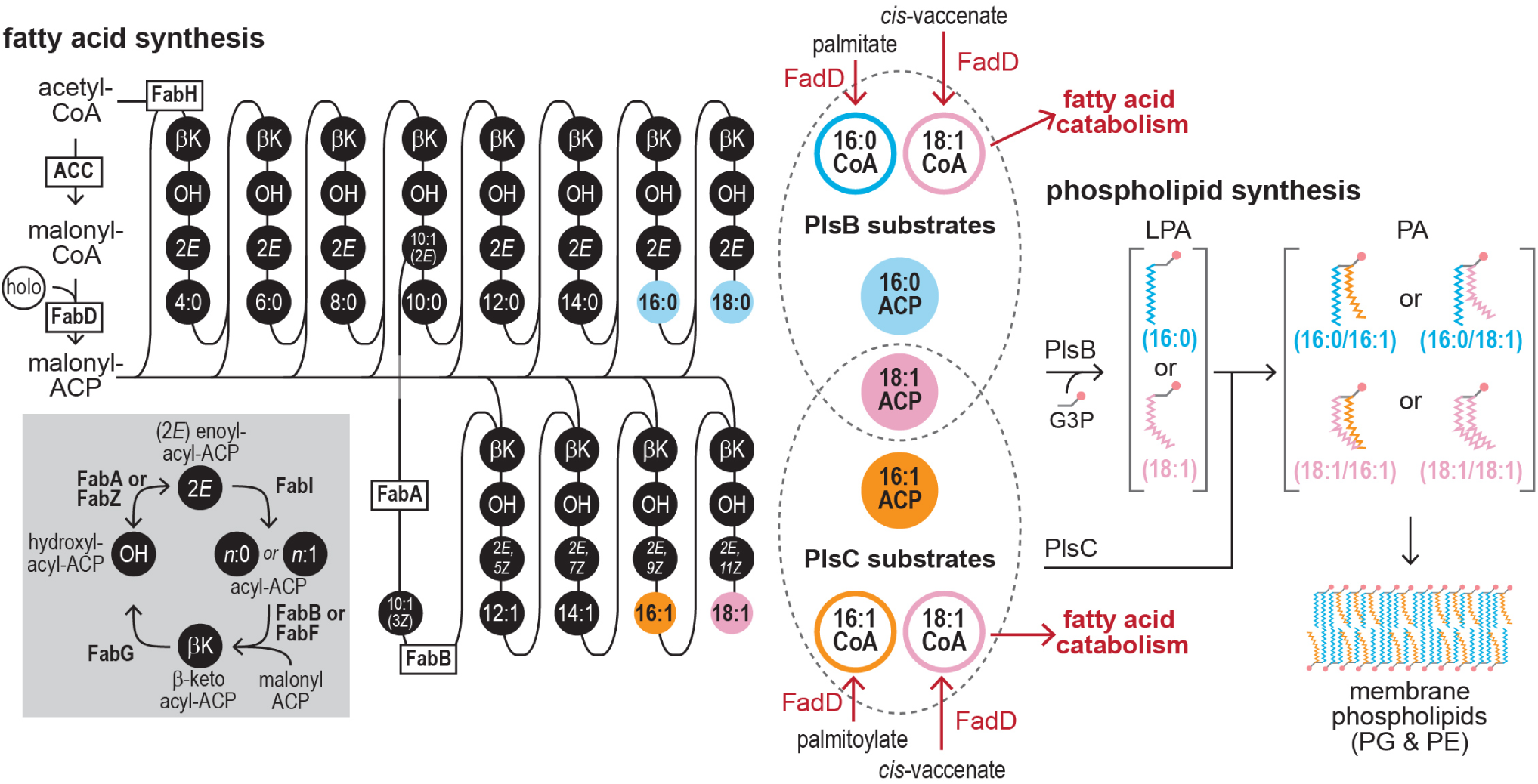
The convergence of the fatty acid synthesis, fatty acid catabolism, and phospholipid synthesis pathways in *E. coli.* Fatty acids are synthesized by repeated cycles of elongation, dehydration, and reduction of acyl-ACP thioesters that ultimately yield long-chain fatty acyl-ACP. The *E. coli* pathway splits into saturated and unsaturated branches at the 10-carbon acyl-ACP intermediate C10:1(2*E*) ACP. FabA reversibly isomerizes (C10:1(2*E*)) with the precursor of the unsaturated pathway (C10:1(3*Z*) ACP). These precursors are irreversibly committed to the saturated and unsaturated branches by FabI and FabB, respectively. Phospholipid synthesis enzymes PlsB and PlsC use long-chain acyl-ACP products of fatty acid synthesis and acyl-CoA intermediates of fatty acid catabolism of appropriate length. PlsB initiates phospholipid synthesis by transferring acyl chains from preferred ACP or CoA substrates (16:0 or 18:1) to the *sn*-1 position of glycerol-3-phosphate (G3P), producing lysophosphatidic acid (LPA). PlsC transfers acyl groups from its preferred ACP or CoA substrates (16:1 or 18:1) to the *sn-*2 position of LPA, generating phosphatidic acid (PA). The relative proportions of acyl chains at *sn*-1 and *sn-*2 positions are determined by the corresponding proportion of acyl thioester substrates in the PlsB and PlsC substrate pools. Note that the substrate preferences of PlsB and PlsC are not absolute: PlsB incorporates a small amount of 16:1 acyl chains at *sn*-1 (37), while PlsC incorporates a small amount of 16:0 acyl chains at *sn-*2. PA is further converted via phospholipid headgroup addition and modification reactions to produce membrane phospholipids phosphatidylglycerol (PG) and phosphatidylethanolamine (PE).

Saturated and unsaturated long-chain fatty acids are also produced by the *E. coli* type II fatty acid synthesis pathway in the form of fatty acyl thioesters attached to an acyl carrier protein (acyl-ACP), which are also substrates for PlsB and PlsC. The ability of PlsB and PlsC to use both acyl-ACP and acyl-CoA as phospholipid precursors places *de novo-*synthesized fatty acids in a common substrate pool with CoA-activated exogenous fatty acids.

Endogenous fatty acid synthesis pathways respond to exogenous fatty acid uptake. In some bacteria, this trait has profound consequences: the ability of *Streptococcus pneumoniae* to inhibit *de novo* synthesis when supplied with exogenous fatty acids allows the bacteria to resist fatty acid synthesis inhibitors (12). Furthermore, as incorporating exogenous fatty acids impacts membrane lipid composition, the composition of the endogenously-generated fatty acid pool must adapt to conserve key membrane properties such as fluidity and thickness. This response is a form of homeoviscous adaptation and is similar to the adaptation of membrane composition to temperature (13). The response to exogenous fatty acids is directed by two transcriptional regulators (FadR and FabR) in *E. coli*. FadR, which is activated by long-chain acyl-CoA, primarily controls expression of the fatty acid degradation pathway (14) and influences expression of several fatty acid synthesis genes (15). The transcriptional regulator FabR controls expression of the fatty acid synthesis enzymes FabA and FabB, which allocate flux between the saturated and unsaturated pathways (16, 17) (**Figure 1**). Exogenous fatty acids may also affect fatty acid synthesis through post-translational mechanisms. In particular, the enoyl-ACP reductase enzyme FabI was reported to be inhibited by acyl-CoA (18), a finding that is often proposed to explain the effects of exogenous fatty acids on membrane assembly and homeostasis (19, 20). The total flux passing through the fatty acid synthesis pathway is also regulated by long-chain acyl-ACP, which inhibit acetyl-CoA carboxylase (ACC) (21) (**Figure 1**). Concentrations of long-chain acyl-ACP (and thus ACC activity) are determined by PlsB activity, which is itself allosterically regulated to coordinate phospholipid synthesis with growth (22). How acyl-CoA might use allosteric or transcriptional mechanisms to regulate fatty acid synthesis is unclear.

Here, we determined the effect of exogenous fatty acids on fatty acid and phospholipid synthesis pathways by comprehensively quantifying fatty acid and phospholipid synthesis intermediates and enzymes. We find that acyl-CoA post-translationally inhibits *de novo* fatty acid synthesis by causing acyl-ACP substrates of PlsB and PlsC to accumulate, which in turn inhibits fatty acid synthesis initiation by ACC. We find no evidence that acyl-CoA inhibits FabI. A mathematical model simulating substrate competition between ACP and CoA substrates closely recapitulates our experimental results. Our findings demonstrate how the cellular demand for phospholipid synthesis retains control over fatty acid synthesis flux regardless of the availability of exogenous fatty acids.

## Results

### Acyl-CoA from exogenous fatty acids immediately inhibits *de novo* fatty acid synthesis

To characterize how the *E. coli* fatty acid pathway responds to the uptake and incorporation of exogenous fatty acids, we quantified acyl-ACP intermediates before and after feeding fatty acids to an exponential-phase culture. Cultures of *E. coli* NCM3722 were prepared at 37 °C in defined minimal medium (MOPS/0.2% glycerol). When culture density reached an optical density of ∼0.4, palmitate (C16:0 fatty acid) was added to achieve final concentration of 0.8 mM. Samples for analysis by liquid chromatography coupled with mass spectrometry (LCMS) were removed from the culture immediately before and after palmitate feeding (23). Surprisingly, concentrations of key acyl-ACP responded within 1 minute after palmitate feeding (**Figure 2A**). The fatty acid initiation and elongation substrate malonyl-ACP decreased by nearly 50%, suggesting inhibited synthesis of malonyl-CoA by ACC. Interestingly, palmitate feeding immediately increased the PlsB substrate C16:0 ACP by ∼2-fold while unsaturated long-chain acyl-ACP C16:1 and C18:1 remained relatively unchanged. Hydroxyl-acyl-ACP and 2(*E*)-enoyl-acyl-ACP species, whose concentrations tightly correlate with fatty acid synthesis flux during steady-state growth (22), also decreased by approximately 50% (**Figure 2B**). Palmitate did not affect acyl-ACP pools of a strain lacking FadD (NCM3722 Δ*fadD*), confirming that effects observed are driven by C16:0 CoA synthesized from exogenous palmitate (**Figure 2A**).

**Figure 2.**
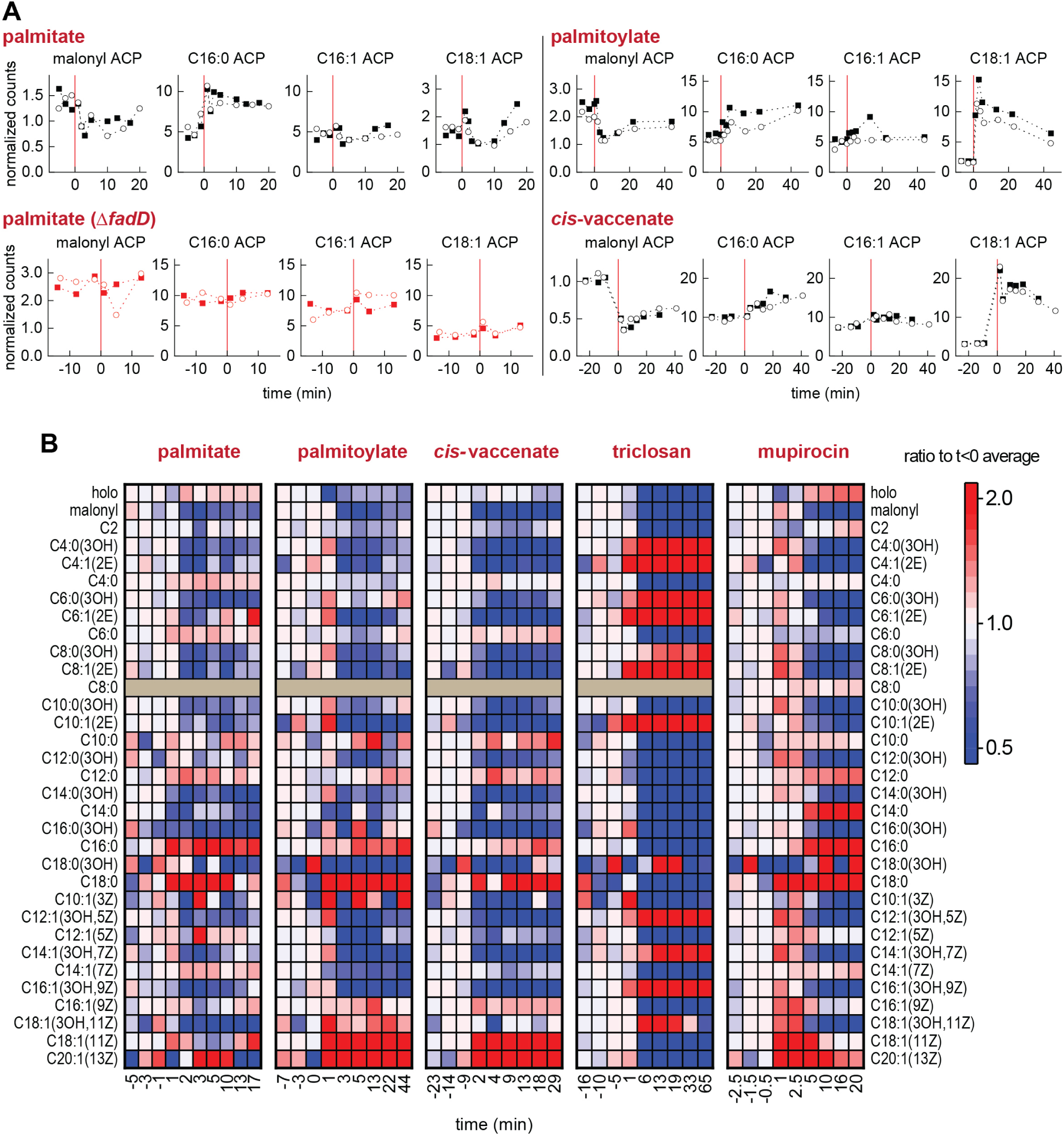
Acyl-ACP pools respond immediately to acyl-CoA synthesized from exogenous fatty acids added to the growth medium. **A.** Concentrations of acyl-ACP quantified before and after addition of indicated exogenous fatty acid (added at 0 min, indicated with vertical red lines). All counts normalized to counts obtained from a non-acylated ACP peptide (LVMALE). Responses of two biological replicates in each condition are displayed; each point represents a single measurement. **B.** Heat map representation of all acyl-ACP pools following acyl-CoA synthesis, triclosan addition, and mupirocin treatment. All values normalized to average of three measurements before treatments. Data are representative of two biological replicates. Triclosan data from (37) and mupirocin data from (22). Measurements of C8:0 ACP lost in some datasets due to recording error.

We next tested the effects of adding unsaturated fatty acids palmitoylate (C16:1(9*Z*)) and *cis*-vaccenate (C18:1(11*Z*)) on the acyl-ACP pools. Similar to palmitate, both fatty acids immediately decreased malonyl-ACP by approximately 50% (**Figure 2A**) and decreased concentrations of hydroxyl-acyl-ACP and 2(*E*)-enoyl-acyl-ACP by ∼50% (**Figure 2B**), again consistent with fatty acid synthesis inhibition. However, unlike palmitate, both unsaturated fatty acids increased C18:1 ACP by ∼5-fold and triggered a relatively small increase in C16:0 ACP. The accumulation of long-chain acyl-ACP and simultaneous reduction of malonyl-ACP and acyl-ACP intermediates that correlate with fatty acid flux strongly suggests that acyl-CoA inhibits fatty acid synthesis by triggering feedback inhibition of ACC by long-chain acyl-ACP. Interestingly, exogenous fatty acids increased the low-abundance C18:0 and C20:1 ACP by severalfold (**Figure 2B**).

### Fatty acid synthesis inhibition by acyl-CoA does not resemble inhibition by the FabI inhibitor triclosan

The response of the acyl-ACP pools to acyl-CoA is far too rapid (∼1 minute) to be mediated by transcriptional regulation. Therefore, acyl-CoA must inhibit fatty acid synthesis via direct interactions with enzymes within either the fatty acid or phospholipid synthesis pathways. C16:0 CoA has been proposed to inhibit fatty acid synthesis by inhibiting the enoyl-acyl ACP reductase FabI (24). This conclusion is based upon *in vitro* experiments which used enoyl-acyl-CoA as a substitute for the actual FabI substrate enoyl-acyl-ACP. However, the effects of acyl-CoA on acyl-ACP pools that we observe do not resemble the effects of the FabI inhibitor triclosan (25). In contrast with the acyl-CoA response, triclosan entirely depleted all long-chain acyl-ACP (**Figure 2A**). While triclosan also depletes malonyl-ACP, concentrations of hydroxyl-acyl ACP and (2*E*)-enoyl-acyl ACP dramatically increase by several fold, a response opposite to the 2-fold depletion of hydroxyl-acyl ACP and (2*E*)- enoyl-acyl ACP caused by acyl-CoA (**Figure 2B**). Furthermore, all acyl-ACP products of FabI decrease, a response not triggered by acyl-CoA. The accumulation of FabI substrates ((2*E*)-enoyl-acyl-ACP, which rapidly equilibrates with hydroxyl-acyl-ACP) and depletion of FabI products is consistent with FabI inhibition. Finally, triclosan depletes holo-ACP, while acyl-CoA does not consistently affect holo-ACP abundance. Overall, the stark differences between the acyl-ACP pool dynamics following acyl-CoA synthesis and triclosan inhibition indicate that acyl-CoA does not suppress fatty acid synthesis *in vivo* by inhibiting FabI.

### Acyl-CoA synthesized from exogenous fatty acids rapidly alters lipid composition of phospholipid intermediates

The effects of acyl-CoA on acyl-ACP pools, in particular the rapid depletion of malonyl-ACP and hydroxyl-acyl-ACP species, resemble the effects of phospholipid synthesis arrest by the alarmone guanosine tetraphosphate (ppGpp) (22) (**Figure 2B**). Inhibition of phospholipid synthesis by ppGpp triggers accumulation of both saturated and unsaturated long-chain acyl-ACP. Long-chain acyl-ACP inhibits synthesis of malonyl-CoA by ACC, which in turn decreases malonyl-ACP concentrations and thus the elongation reactions catalysed by β-keto-acyl-ACP synthases FabH, FabB, and FabF. This is evident from decreased concentrations of malonyl-ACP as well as depletion of flux-sensitive intermediates hydroxyl-acyl ACP and (2*E*)-enoyl-acyl ACP when ppGpp is elevated (22).

Furthermore, phospholipid synthesis inhibition by ppGpp also depletes the phospholipid synthesis intermediates PA and phosphatidylserine (PS). However, unlike phospholipid synthesis inhibition by ppGpp, acyl-CoA synthesized from exogenous fatty acids does not substantially deplete PA and PS (**Figure 3A**). Instead, within 1-2 minutes of feeding, the acyl chain compositions of phospholipid synthesis intermediates immediately change (**Figure 3B**). Specifically, palmitate slightly increases the fraction of PA bearing 16:0 acyl chains at *sn-*1, consistent with greater incorporation of C16:0 fatty acid by PlsB from C16:0 CoA. Strikingly, palmitate also increases the proportion of phospholipids with C16:0 at *sn-*2, consistent with utilization of C16:0 CoA by PlsC. These changes in the composition of the PA intermediate pool propagate over time to alter the lipid composition of the membrane phospholipids PG and PE (**Figure 3B**).

**Figure 3.**
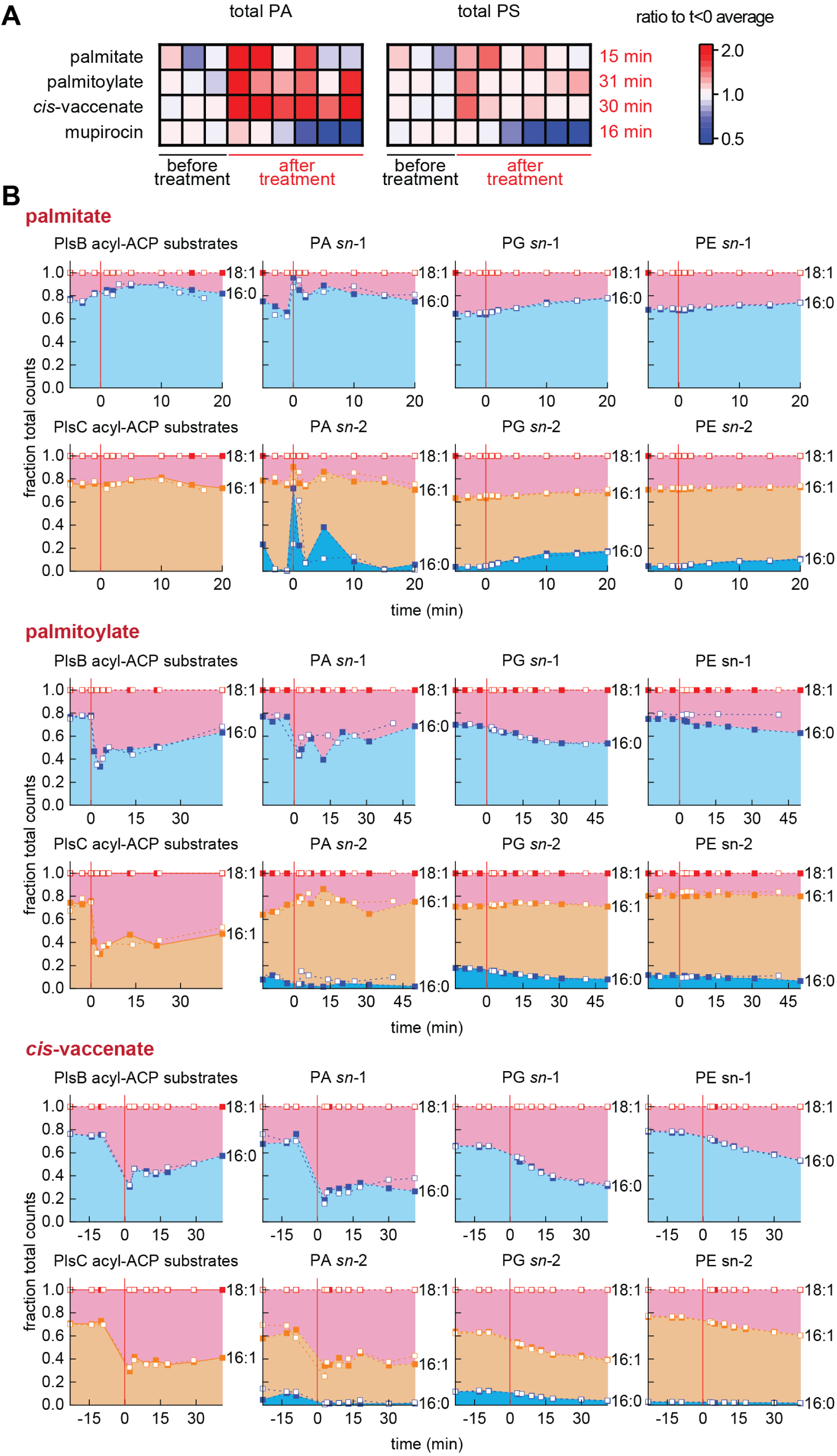
Phospholipid intermediate and membrane phospholipid pools respond immediately to acyl-CoA synthesis. **A.** Relative abundance of PA and phosphatidylserine (PS) immediately following exogenous fatty acid feeding compared with phospholipid synthesis inhibition by ppGpp (ppGpp synthesis triggered by mupirocin). All values normalized to average of three measurements before treatments. Data are representative of two biological replicates. Mupirocin data from (22). Values on right axis indicate time of final measurement shown. **B.** Comparison of PlsB and PlsC substrate pools with composition of PA and membrane phospholipids PG and PE before and after addition of exogenous fatty acids. Areas represent fraction of total LCMS counts for phospholipid species bearing indicated acyl chains at either the *sn-*1 or *sn-*2 positions. All kinetic series obtained from 2 independent experiments, which are distinguished by closed and open symbols.

Addition of the unsaturated fatty acid palmitoylate also rapidly changed the composition of the PA intermediate pool. Although C16:1 CoA is not expected to significantly contribute to the PlsB substrate pool, palmitoylate immediately increased the proportion of C18:1 *sn-*1 PA (**Figure 3B**).

This is due to the sudden increase in C18:1 ACP caused by C16:1 CoA. Similarly, palmitoylate rapidly perturbed the composition of the PlsC acyl-ACP substrate pool, with C18:1 ACP becoming the predominant ACP thioester. However, the proportion of lipids incorporated at PA *sn-*2 did not match the proportions represented within the PlsC acyl-ACP substrate pool, reflecting the presence of C16:1 CoA countering the increased concentration of C18:1 ACP. *Cis-*vaccenate feeding dramatically increased the proportion of C18:1 ACP within the acyl-ACP pools of both PlsB and PlsC and immediately changed the lipid composition at both the *sn-*1 and *sn-*2 positions of PA, which propagated to the composition of membrane phospholipids PG and PE (**Figure 3B**). These results indicate that exogenous fatty acids alter the lipid composition both directly (through use of acyl-CoA substrates by PlsB and PlsC) and indirectly (by changing the composition of the long-chain acyl-ACP pool). Importantly, the indirect effects on the acyl-ACP pools further increase the incorporation of saturated or unsaturated fatty acids from the acyl-ACP pools into the membrane, thus amplifying the influence of acyl-CoA on the balance of saturated/unsaturated membrane lipids.

### A mathematical model simulating competition between acyl-ACP and acyl-CoA for PlsB and PlsC reproduces acyl-ACP perturbations and fatty acid synthesis inhibition

While it is clear how acyl-CoA is able to rapidly change the lipid composition of newly-synthesized phospholipids, it is less obvious how acyl-CoA triggers acyl-ACP accumulation and fatty acid synthesis inhibition. The strong and rapid effects of acyl-CoA on the lipids incorporated into phospholipids indicate that acyl-ACP and acyl-CoA compete for PlsB and PlsC active sites. This competition in turn suggests a mechanism by which acyl-CoA decreases malonyl-ACP concentrations and thereby attenuates fatty acid flux: as a competing substrate, acyl-CoA decreases acyl-ACP consumption by PlsB and PlsC, causing long-chain acyl-ACP to accumulate and inhibit malonyl-CoA synthesis by ACC. This is consistent with the decreased concentrations of malonyl-ACP and other flux-sensitive acyl-ACP (hydroxyl-acyl-ACP and (2*E*)-enoyl-acyl-ACP).

To test this notion we constructed a mathematical model of the *E. coli* fatty acid and phospholipid synthesis pathways using the simulation software COPASI (26). For simplicity, the simulated fatty acid synthesis pathway included only initiation and elongation steps. Fatty acid synthesis initiation reactions were combined to a single ACC-FabD reaction generating malonyl-ACP from holo-ACP and acetyl-CoA, followed by FabH condensing malonyl-ACP with acetyl-CoA (**Figure 4A**). All subsequent elongation reactions were represented by a “FabB” reaction modified to account for combined chain length preferences of β-keto-acyl-ACP synthases FabB and FabF (27). The branch point between saturated and unsaturated pathways was simulated as two parallel reactions consuming C10:0 ACP as a common substrate. A term implementing feedback inhibition of the ACC-FabD reaction by the long-chain acyl-ACP pool was also included. All enzymatic reactions were simulated as one- or two-substrate reactions using Michaelis-Menten kinetics with the exception of the reactions catalysed by PlsB and PlsC. To simulate substrate competition, the separate binding/unbinding and catalytic steps of acyl-ACP and acyl-CoA substrates with PlsB and PlsC were explicitly included (**Figure 4B**), following the example of recent modelling work (28). In order to maintain overall phospholipid flux at a fixed value despite the changing abundance of PlsB and PlsC substrates, a feedback inhibition term controlling PlsB activity based on total phospholipid content was included. Generation of acyl-CoA was simulated by a stepwise change in acyl-CoA concentration after the simulation had reached steady-state. Further details of the simulation are included within the Supplementary Information.

**Figure 4.**
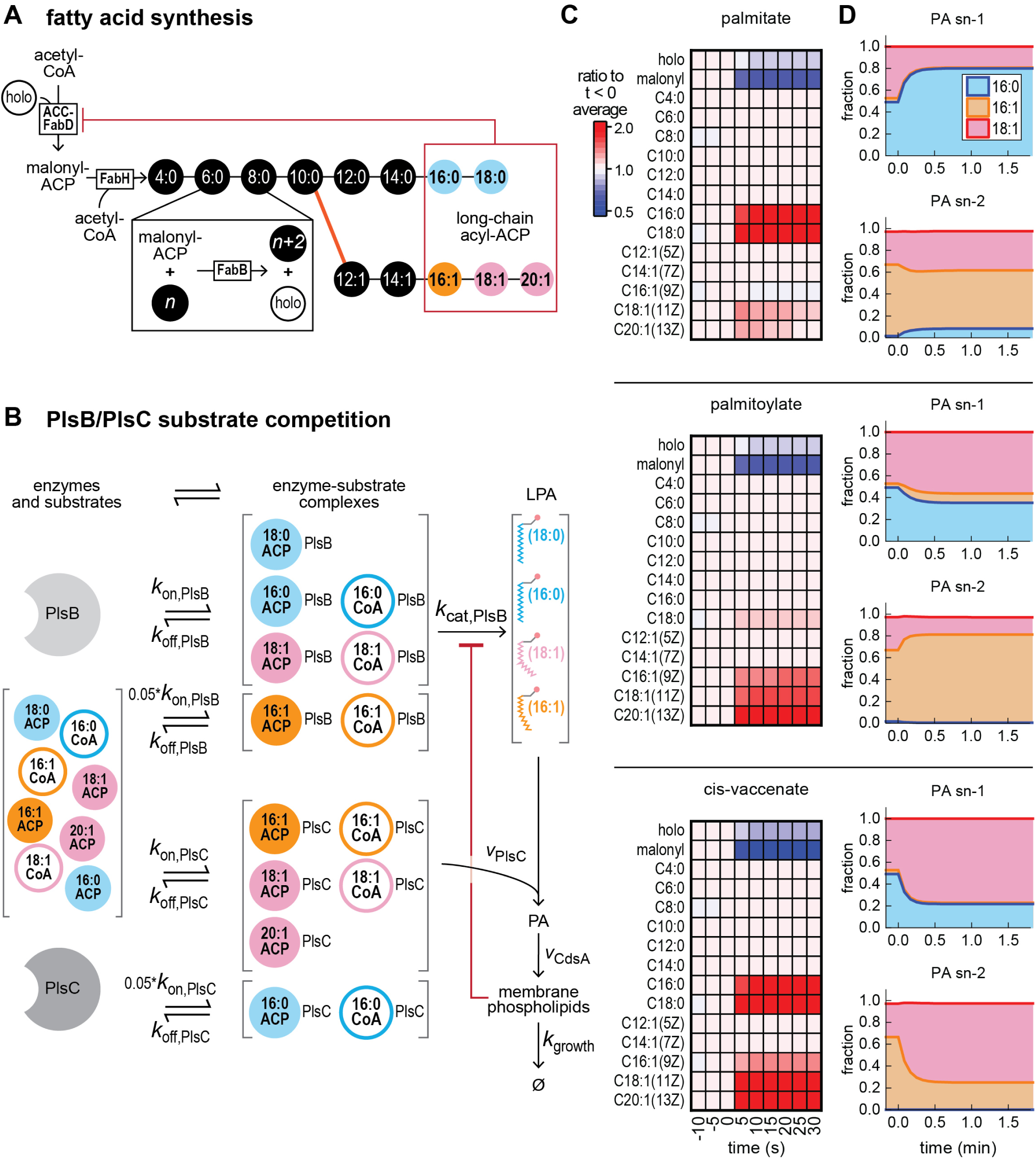
A mathematical model of the fatty acid and phospholipid synthesis pathways that explicitly simulates substrate competition between acyl-CoA and acyl-ACP reproduces experimental results. **A, B.** Diagram of the simplified fatty acid pathway **(A)** and substrate competition scheme **(B)** used in the simulation. Substrate preferences for PlsB (disfavouring C16:1 ACP and C16:1 CoA) and PlsC (disfavouring C16:0 ACP and C16:0 CoA) are implemented by decreasing the binding rates (*k*_on_) for these substrates. **C, D.** Simulation output for acyl-ACP pools **(C)** and phospholipid intermediate lipid compositions **(D)** after exogenous fatty acid feeding at 0 min. Detailed explanation of the model, including differential equations and parameters used, are included in the Supplementary Information.

Similar to our experimental observations, the simulated stepwise increase in C16:0 CoA (emulating palmitate feeding) immediately increased both C16:0 and C18:0 ACP and decreased malonyl-ACP (**Figure 4C**). The increase in C16:0 ACP is the direct result of substrate competition between C16:0 ACP and C16:0 CoA, while the increase in C18:0 ACP results from increased abundance of substrate for the elongation reaction to 18:0 ACP. Furthermore, our model also reproduced the changing composition of the PA pool, with C16:0 increasing at *sn*-1 and *sn-*2 (**Figure 4D**). Simulated production of C16:1 CoA (equivalent to palmitoylate feeding) also closely followed our experimental observations. C16:1 CoA increased long-chain unsaturated acyl-ACP and decreased malonyl-ACP. The increase in C18:1 ACP also increased the fraction of 18:1 *sn*-1 PA. However, just as in our observations, palmitoylate slightly decreased the 18:1 *sn-*2 PA pool, reflecting the incorporation of palmitoylate from C16:1 CoA by PlsC. Finally, simulated responses of the acyl-ACP pools to *cis-*vaccenate feeding also closely followed our experimental observations: both saturated and unsaturated long-chain acyl-ACP increased significantly while malonyl-ACP decreased. The increased proportions of 18:1 at both *sn-*1 and *sn-*2 PA after *cis-*vaccenate feeding also mirror our experimental observations.

### The FabA/FabB ratio is adjusted to restore proportions of saturated and unsaturated acyl-ACP within the PlsB substrate pool

The changes in membrane composition induced by exogenous fatty acid incorporation affect biophysical properties of the phospholipid membrane such as membrane fluidity and thickness. To maintain the biophysical properties of its membrane, *E. coli* controls relative synthesis rates of saturated and unsaturated fatty acids by adjusting expression of the branch point enzymes FabA and FabB: decreasing the FabA/FabB ratio increases the production of unsaturated fatty acids, while increasing FabA/FabB increases the production of saturated fatty acids (17). We quantified proteins of the fatty acid synthesis pathway using LCMS to determine the transcriptional response to exogenous fatty acids. After palmitate feeding, FabB steadily increased and reached final steady-state level after approximately 2 hours. In contrast, FabA levels remained constant (**Figure 5A**), causing the FabA/FabB ratio to decrease (**Figure 5B**). These changes were not observed in cultures treated with tergitol or in cultures of palmitate-fed Δ*fadD*. In contrast, addition of unsaturated fatty acids palmitoylate and *cis-*vaccenate decreased both FabA and FabB. After two generation periods, the FabA/FabB steady-state ratios increased slightly, consistent with the expected homeostatic response.

**Figure 5.**
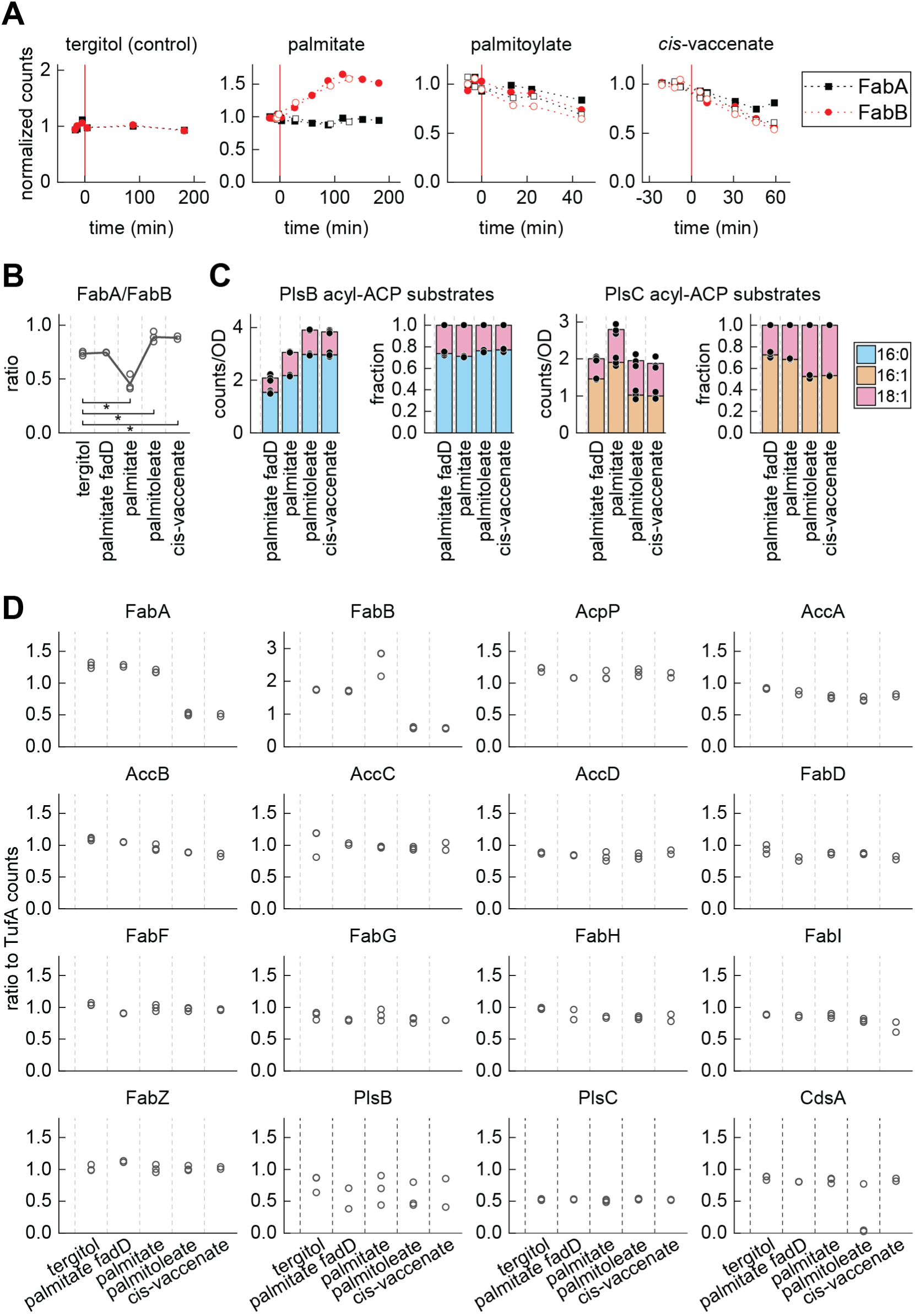
Transcriptional response of the fatty acid pathway to exogenous fatty acids. **A.** Normalized response (*t*<0 average = 1) of FabA and FabB concentrations after feeding of exogenous fatty acids compared with tergitol control. Data from two biological replicates displayed, which are distinguished by closed and open symbols. **B, C, D.** Steady-state values of FabA/FabB ratio (**B**), total LCMS counts and fractions of total counts of PlsB and PlsC acyl-ACP substrates (**C**), and concentrations of fatty acid synthesis pathway enzymes (**D**) measured two doublings after feeding exogenous fatty acids. For all plots, symbols represent multiple measurements from one culture; line (**B**) or bars (**C**) represent average value. For statistical comparison in **B**, asterisk indicates significant differences between fatty acid feeding conditions and tergitol control with *p* <0.02 (Student’s *t*-test, 2-tailed unequal distribution performed using Microsoft Excel T.TEST function). For measurements of fatty acid synthesis pathway enzymes at steady-state in tergitol (**D**), palmitate-fed wild-type culture, and palmitoleic acid-fed culture, three measurements were performed. For *cis-*vaccinate and palmitate-fed Δ*fadD* cultures two measurements were performed.

The changes in the FabA/FabB ratio adapts the abundance of long-chain acyl-ACP and the relative proportions within the PlsB and PlsC substrate pools (**Figure 5C**). After two doublings in palmitate-supplemented medium, concentrations of unsaturated acyl-ACP increase by approximately 50%, whereas C16:0 ACP increases after two doublings in medium supplemented with either palmitoylate or *cis*-vaccenate. Surprisingly, despite these adaptations in the fatty acid synthesis pathway, the final steady-state fraction of C16:0 ACP within the PlsB substrate pool in all three fatty acid-supplemented conditions matched the ratio existing before fatty acid addition (∼0.8). This suggests that the transcriptional response to exogenous fatty acids restores the C16:0/C18:1 ACP ratio existing prior to exogenous fatty acid feeding. We also quantified the remaining enzymes of the fatty acid synthesis pathway during steady-state growth in the presence of exogenous fatty acids. Interestingly, with the exception of FabA and FabB, all enzyme concentrations remained constant, as did the first three enzymes of phospholipid synthesis (PlsB, PlsC, CdsA) (**Figure 5D**). This indicates that expression of the fatty acid synthesis pathway remains constant despite the availability of exogenous fatty acids. Thus, while transcriptional regulation adjusts the relative proportions of saturated and unsaturated acyl-ACP, it does not downregulate expression of the fatty acid synthesis pathway.

## Discussion

The conversion of exogenous fatty acids to substrates for membrane synthesis inhibits endogenous fatty acid synthesis. Substrate competition between acyl-ACP and acyl-CoA triggers the same feedback mechanism that coordinates fatty acid synthesis with phospholipid synthesis and cell growth: accumulated long-chain acyl-ACP attenuate malonyl-CoA synthesis by ACC. This ensures that fatty acid synthesis is regulated by PlsB activity regardless of the presence of exogenous fatty acids. We find no evidence that acyl-CoA inhibits FabI *in vivo.* We also find that the transcriptional response to exogenous fatty acids is limited to restoring the balance between saturated and unsaturated fatty acid synthesis by tuning FabA and FabB, as the concentrations of all other fatty acid synthesis pathway enzymes remain unchanged. This reflects the requirement for intermediates of the fatty acid synthesis pathway (including the lipopolysaccharide precursor C14:0-OH ACP) and ensures that fatty acid synthesis resumes rapidly when the supply of exogenous fatty acids is exhausted. This design takes advantage of the superior reversibility and responsiveness of post-translational regulation compared to transcription-driven responses (22). Our finding is important for understanding the potential signalling role of acyl-CoA generated from the breakdown of mislocalized phospholipids in the outer leaflet of the outer membrane (20, 29).

We also find that exogenous fatty acids perturb the membrane composition not only through direct incorporation, but also by changing the ratio of saturated and unsaturated long-chain acyl-ACP. This amplifies the effects of exogenous fatty acids on membrane composition: palmitate increases the abundance of C16:0 ACP relative to C18:1 ACP, while palmitoylate and *cis-*vaccenate increase C18:1 ACP relative to C16:0 ACP. A transcriptional response corrects for this effect by altering the FabA/FabB ratio. The transcriptional regulator FabR binds both C16:0 and C18:1 ACP, with the FabR-C18:1 ACP complex repressing *fabB* transcription and thereby establishing a negative feedback loop that attenuates unsaturated fatty acid synthesis (16). Previously, the observation that FabR adjusts FabA and FabB expression in response to acyl-CoA led to the proposal that FabR is able to bind both acyl-ACP and acyl-CoA. However, we find that acyl-CoA directly changes the C16:0/C18:1 ACP ratio due to substrate competition, with saturated acyl-CoA increasing C16:0 ACP and unsaturated acyl-CoA increasing C18:1 ACP. Therefore, it is possible that FabR monitors only the C16:0/C18:1 ACP ratio and does not bind acyl-CoA *in vivo*. A close examination of the data in Zhu *et al.* suggests that FabR exhibits far higher affinity to acyl-ACP than acyl-CoA (16). *In vitro* competition studies between acyl-CoA and acyl-ACP for triggering DNA binding by FabR could address this question.

Could the substrate competition-driven inhibition of fatty acid synthesis we observe here occur in other bacteria? The enzyme PlsB, which uses both acyl-CoA and acyl-ACP, occurs rarely in bacterial species. Most bacterial species initiate phospholipid synthesis using PlsX and PlsY, which generate LPA in two steps: PlsX converts acyl-ACP to acyl-phosphate, which is subsequently converted to LPA by the G3P acyl transferase PlsY. PlsX and PlsY cannot use acyl-CoA as substrates for the acyltransferase reaction (30). However, many bacteria use a fatty acid kinase (FakA) to convert exogenous fatty acids to acyl-phosphate (31). As the reaction catalysed by PlsX is reversible, acyl-phosphate generated from exogenous fatty acids may be converted to acyl-ACP, which can be subsequently fed into the fatty acid synthesis pathway and elongated to long-chain acyl-ACP (**Figure 6**). Therefore, as the uptake of fatty acids via FakA and PlsX may also lead to acyl-ACP accumulation, PlsB may not be necessary for exogenous fatty acids to inhibit fatty acid synthesis. The use of acyl-ACP synthases by organisms such as *Chlamydia trachomatis* and *Neisseria gonorrhoeae*, which synthesize acyl-ACP from exogenous fatty acids in a single step, suggests a more straightforward pathway for generating long-chain acyl-ACP that could inhibit *de novo* fatty acid synthesis (1, 32). Thus, all exogenous fatty acid incorporation pathways characterized to date contribute to a substrate pool that is shared with products of fatty acid synthesis. If the inhibition of fatty acid synthesis by long-chain acyl-ACP or acyl-phosphates (at ACC or another step) is also well-conserved, then exogenous fatty acids may inhibit endogenous fatty acid synthesis by substrate competition in many other species.

**Figure 6.**
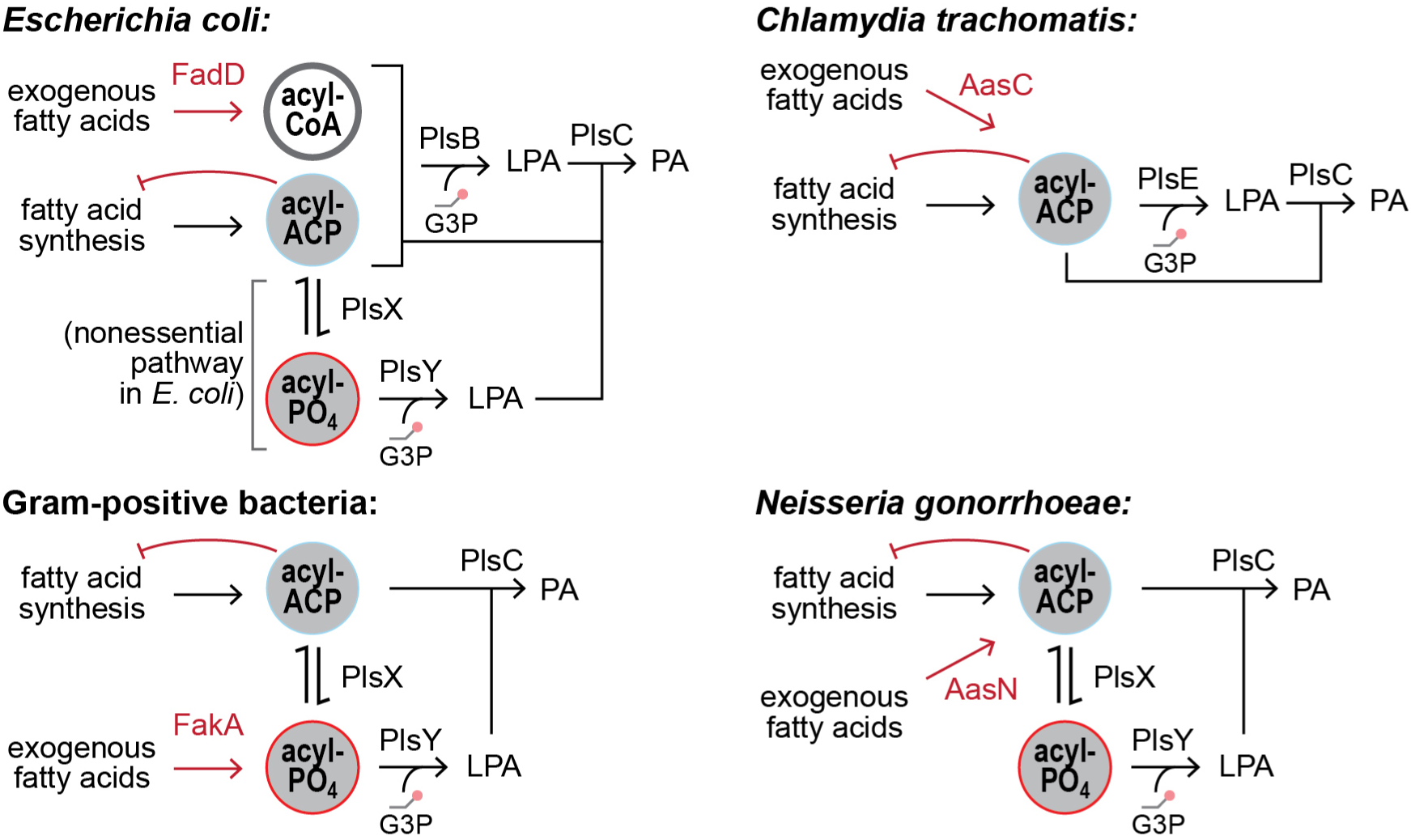
The conserved intersections of fatty acid synthesis, exogenous fatty acid incorporation, and phospholipid synthesis pathways potentially enables exogenous fatty acids to inhibit *de novo* fatty acid synthesis in diverse bacteria.

The inhibition of fatty acid synthesis by exogenous fatty acids is highly relevant for determining whether an organism can use exogenous fatty acids to evade the effects of fatty acid synthesis-targeting antibiotics. Exogenous fatty acids strongly inhibit malonyl-CoA synthesis by ACC in *Streptococcus pneumoniae,* although the mechanism was not clear (12). This inhibition proved crucial for preserving sufficient holo-ACP despite treatment with FabF and FabI inhibitors (which deplete holo-ACP) to allow exogenous fatty acid incorporation. In contrast, exogenous fatty acids could not sufficiently inhibit malonyl-CoA synthesis by ACC in *Staphylococcus aureus*. As a consequence, treatment with FabF and FabI inhibitors entirely depletes holo-ACP due to continued synthesis of malonyl-ACP (as we observe in *E. coli*). Depletion of holo-ACP prevents exogenous fatty acid incorporation and arrests growth. Investigating how exogenous fatty acids stringently inhibit ACC in *S. pneumoniae* would provide important insight into an antibiotic resistance mechanism.

## Methods

### Strains, chemicals, and growth conditions

*Escherichia coli* K-12 strain NCM3722 (CGSC #12355) was used for all experiments. NCM3722 Δ*fadD* was constructed using lambda-red recombination (33) with primers listed in **Table 1**. Cultures were prepared in Erlenmeyer flasks with 0.2% w/vol glycerol in MOPS minimal medium (34). Flasks were maintained at 37 °C by submersion in a Grant Instruments Sub Aqua Pro water bath and stirred (1200 rpm) using a magnetic stir bar (20 mm) coupled to a stir plate (2mag MIXdrive 1 Eco and MIXcontrol 20). Growth was monitored with optical density measurements (Ultrospec 10 Cell Density Meter, GE Healthcare). Fatty acids (40 mM) were neutralized to pH 7 using NaOH and solubilized with 26% tergitol before dilution in culture medium. Fatty acids were obtained from Sigma-Aldrich: palmitic acid (P0500), palmitoleic acid (P9417) and, *cis-*vaccenic acid (V0384).

**Table 1.**
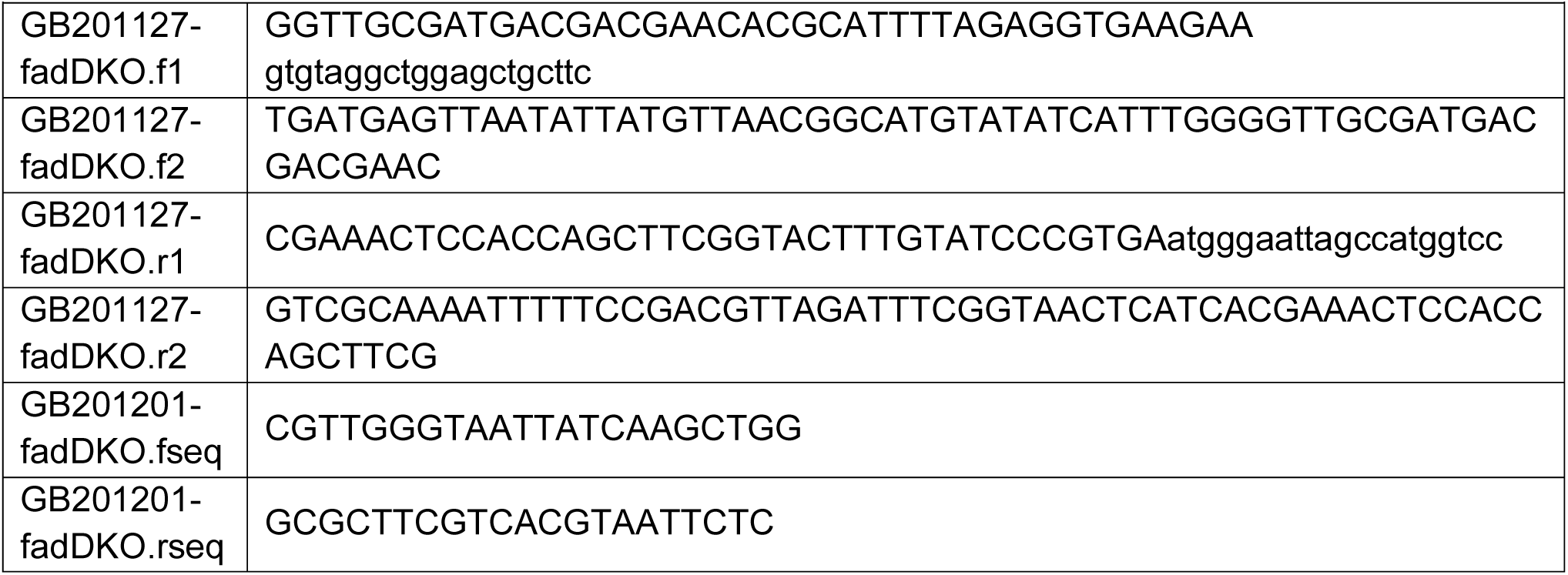
Primers used for constructing and confirming ΔfadD strain.

Unsaturated fatty acids were stored at -20 °C and prepared as solutions immediately before use.

### Culture sampling for LCMS analysis

Samples (usually 1 mL) for all analysis modalities were rapidly removed from cultures and quenched by pipetting directly into ice-cold 10% trichloroacetic acid (TCA) to 2% final TCA concentration, pelleted, and stored at -80 °C until processing for analysis.

### LCMS quantification and analysis

LCMS methods are fully described in reference (22). LCMS was performed using Agilent LCMS (binary pump (G1312B), autosampler (G7167A), temperature-controlled column compartment (G1316A), and triple quadrupole (QQQ) mass spectrometer (G6460C) equipped with a standard ESI source) all operated using MassHunter (version 7.0). Mass spectrometer set in dynamic MRM mode using transitions generated *in silico* by a script written in Python using RDkit library.

Transitions for targeted proteomics assays were developed using Skyline (35). LC/MS data was further processed in Skyline versions 4.x using an *in silico* generated transition list for targets and corresponding internal standards.

#### Acyl-ACP quantification

Sample pellets were lysed by suspending quenched pellets in a buffered urea solution (10 mL of 50 mM potassium phosphate buffer, pH 7.2, 6 M urea, 10 mM N-ethyl-maleimide, 5 mM EDTA and 1 mM ascorbic acid) with ^15^N-labelled internal standards (generated using U-^15^N *E. coli* whole cell extracts from MOPS minimal medium cultures with ^15^NH_4_Cl as the sole nitrogen source) and proteins precipitated according to (22). Protein pellet was resolubilized in 10 μL digestion buffer (4% 2-octyl-glucoside, 25 mM potassium phosphate pH 7.2) and digested initiated by adding 10 μL of 0.1 mg/mL GluC protease (Promega). Digestion continued overnight at 37 °C. After quenching with 5 μL MeOH, samples were centrifuged and 10 μL injected into LC/MS. Separation was performed on 2.1 mm x 50 mm 1.7 μm CSH C-18 column (Waters) held at 80°C using a binary gradient: 15% B, 3 minute ramp to 25%, 9 min increase to 95% and 1 minute hold at 95% B before 3 minute re-equilibration at starting conditions (A: 25 mM formic acid, B: 50 mM formic acid) at a flow rate of 0.6 mL/min. MS transitions for both ^14^N and ^15^N acyl-ACP were developed as described in (23) and (22).

#### Phospholipid quantification

TCA-quenched sample pellets were resuspended in 150 μL of MeOH, 250 μL of U-^13^C *E. coli* phospholipid extract and 250 μL *m-*tert-butyl ether (MTBE), and subsequently vortexed and sonicated in a water bath. 125 μL of 15 mM citric acid/ 20 mM dipotassium phosphate buffer was added to homogenized pellets and vortexed. Liquid phases were separated by centrifugation for 10 min at 20000g. 450 μL of the upper phase was moved to a new tube and dried in a vacuum centrifuge. Dried lipids were resuspended in 10 μL 65:30:5 (v/v/v) isopropanol/acetonitrile/H2O supplemented with 10 mM acetylacetone. After addition of 5 μL H2O, 5 μL of resulting mixture was injected into the LC/MS system. Separation was performed on 2.1 mm x 50 mm 1.7 μm CSH C-18 column (Waters) at 60°C with a flow rate of 0.6 mL/min using the following binary gradient: 25% B, ramp to 56%B in 6 min followed by linear increase to 80% B in 6 min, 2 min hold at 100% B and 3 min re-equilibration (A: 0.05% NH4OH in water, B: 0.05% NH4OH in 80% isopropanol 20% ACN).

#### Targeted protein quantification

TCA-quenched sample pellets were resuspended in 1 mL of 50 mM potassium phosphate buffer, pH 7.2 and 6 M urea that had been pre-mixed with U-^15^N-labeled *E. coli* internal standards. After precipitation in chloroform/methanol, protein pellets were resuspended in 10 μL of digestion buffer (4% 2-octyl-glucoside in 25 mM Tris buffer pH 8.1, 1 mM CaCl_2_, 5 mM TCEP). Cysteines were alkylated by adding 3 μL of 50 mM iodoacetamide followed by 15 minutes of incubation. Digestion initiated by addition of 10 μL of 0.2 mg/mL Trypsin Gold (Promega). After overnight digestion at 37°C, 10 μL of digestion reaction was injected in LCMS system and separation performed on 2.1 mm x 50 mm 1.7 μm CSH C-18 column (Waters) held at 40°C using a binary gradient: 2% B, 20 minute ramp to 25% B, 4 min increase to 40% B, 0.5 ramp to 80% and 1 minute hold at 80% B before 3 minute re-equilibration at starting conditions (A: 25 mL formic acid, B: 50 mM formic acid) at a flow rate of 0.5 mL/min. Analysis of kinetic data revealed that tergitol addition affected detected protein concentrations. All protein quantities were therefore normalized to the abundance of elongation factor TufA to account for effects of tergitol.

### Mathematical modelling

The mathematical model was constructed using COPASI version 4.39 (build 272). Full details of the model are included in the Supplementary Information.

## Data Availability

The COPASI files (.cps) for the model and all source data for figures will be made available via Figshare at 10.6084/m9.figshare.27302661. The mass spectrometry proteomics data (and acyl-ACP and phospholipids data) have been deposited to the ProteomeXchange Consortium via the PRIDE (36) partner repository with the dataset identifier PXD057199 and will be made available upon publication.

## Acknowledgements

We thank Nicole Scherer for preliminary LCMS analysis and members of the Bokinsky Lab for feedback and scientific discussions. Project supported by start-up funds to GB from the TU Delft Bionanoscience Department.

## Author Contributions

SvdB: designed and performed experiments, built data analysis pipeline, evaluated data. FY, AZ-D: LCMS analysis. GB: supervised project, performed pilot experiments, constructed COPASI model, evaluated data, and wrote the manuscript. All authors approved the final manuscript.

## Competing Interests

The authors declare no competing interests.

## Notes

### Competing Interest Statement

The authors have declared no competing interest.

