## Supplementary Information for "Exogenous fatty acids inhibit fatty acid synthesis through competition between endogenously- and exogenously-generated substrates for phospholipid synthesis in *Escherichia coli*"

### Model description

#### COPASI settings.

COPASI version 4.39 was used for all simulations. The model topology is depicted in Figure 4 in the main text. All data were generated using the "Time Course" Task, simulating 15000 seconds in 5 second intervals with deterministic (LSODA) method with default parameters (relative tolerance $10^{-6}$, absolute tolerance $10^{-12}$, 100000 maximum internal steps, 0 maximum internal step size). Concentrations of all species (except species listed in **Table S1**) were initialized at 0 μM. All concentrations were subsequently allowed to evolve as the simulation progressed unless fixed at set values (indicated in **Table S1**). The simulation was initiated with acyl-CoA set at zero to allow pools of acyl-ACP, phospholipid synthesis intermediates, and phospholipid abundance to reach steady-state from initial conditions. At 8000 seconds, the "Events" function triggered a stepwise increase of acyl-CoA concentration from 0 to 5 μM.

**Table S1.** Initial concentrations of species set at non-zero values.

| | | |
|---|---|---|
| Acetyl-CoA | 600 μM | fixed |
| Holo-ACP | 50 μM | Consumed/regenerated by reactions |
| FabH | 1.6 μM | fixed |
| FabB | 5 μM | fixed |
| PlsB | 0.45 μM | Consumed /regenerated by substrate binding and reactions |
| PlsC | 0.4 μM | Consumed /regenerated by substrate binding and reactions |
| CdsA | 0.2 μM | fixed |
| Acyl-CoA | 0 μM | Fixed, then stepwise change to fixed value (5 μM) to simulate feeding from a large pool of exogenous fatty acids |

Parameters for all equations described below are given in **Table S2.**

#### Initiation (ACC-FabD)

The rate of fatty acid synthesis initiation from acetyl-CoA $v_{\text{ACC}}$ was modelled using a combined ACC-FadD reaction that generates malonyl-ACP from holo-ACP and acetyl-CoA. The rate law used was based on an irreversible bimolecular reaction and included an empirical inhibition term **(Equation S1)**. The concentration of the inhibitor pool (long-chain acyl-ACP) was defined as the sum of free C16:0, C18:0, C16:1, C18:1, and C20:1 ACP concentrations. Acyl-ACP bound within an enzyme-substrate complex with PlsB or PlsC was not included in this term.

**(Eq. S1)**

$$v_{\text{ACC}} = \frac{Vmax*[\text{acetyl-CoA}]*[\text{holo-ACP}]}{Kma*Kmb+[\text{acetyl-CoA}]*Kmb+[\text{holo-ACP}]*Kma+[\text{acetyl-CoA}]*[\text{holo-ACP}]} * \left(\frac{Ki}{Ki+[\text{long chain acyl-ACP}]}\right)$$

#### Initiation (FabH)

The FabH reaction was modelled using a simple irreversible bimolecular rate law ($v_{\text{FabH}}$) that generates C4:0 ACP from acetyl-CoA and malonyl-ACP **(Equation S2)**.

**(Eq. S2)** $$v_{\text{FabH}} = \frac{k_{cat}*[\text{FabH}]*[\text{acetyl-CoA}]*[\text{malonyl-ACP}]}{Kma*Kmb+[\text{acetyl-CoA}]*Kmb+[\text{malonyl-ACP}]*Kma+[\text{acetyl-CoA}]*[\text{malonyl-ACP}]}$$

#### Acyl-ACP elongation ("FabB")

All elongation reactions after FabH were labelled "FabB" (combining FabB and FabF reactions) and were modelled using a simple irreversible bimolecular rate law $v_{\text{FabB}}$ that uses malonyl-ACP and acyl-ACP of length *n* to generate acyl-ACP of length *n*+2 **(Equation S3)**. To account for the different activities of FabB and FabF on acyl-ACP chains of different length and saturation, a coefficient *c*(*n*) was applied for each chain length. These coefficients were chosen based on experimentally-determined chain length-dependent activities of FabB and FabF (1).

For the branch point between saturated and unsaturated synthesis, two parallel reactions consuming C10:0 ACP were implemented that produced either C12:0 or C12:1 ACP.

**(Eq. S3)** $$v_{\text{FabB}} = \frac{c(n)*k_{cat}*[\text{FabB}]*[\text{acyl-ACP}(n)]*[\text{malonyl-ACP}]}{Kma*Kmb+[\text{acyl-ACP}]*Kmb+[\text{malonyl-ACP}]*Kma+[\text{acyl-ACP}]*[\text{malonyl-ACP}]}$$

#### Substrate competition for PlsB and LPA synthesis.

To simulate substrate competition between long-chain acyl-CoA and acyl-ACP, all binding and release steps involving acyl-thioester substrates with PlsB were explicitly included as separate reactions **(Scheme S1)**. This approach takes inspiration from a complete model of the *E. coli* fatty acid synthesis pathway (2). All PlsB-substrate complexes reacted at identical rates, generating free PlsB, the corresponding LPA depending on the acyl chain transferred, and holo-ACP if an acyl-ACP thioester was used as a substrate (free CoA concentrations were not included in the model, so CoA generation from reactions with acyl-CoA is not accounted for). The PlsB substrate glycerol-3-phosphate (G3P) was also not explicitly included (equivalent to setting G3P at a fixed concentration).

$$\text{acyl-thioester + PlsB} \underset{k_{\text{off}}}{\overset{k_{\text{on}}}{\rightleftharpoons}} \text{PlsB-substrate complex} \xrightarrow{k_{\text{cat}}*f([\text{total PL}])} \text{PlsB + LPA (+ holo-ACP)}$$

**(Scheme S1)**

In *E. coli,* the phospholipid/protein ratio is stringently maintained. This means that neither phospholipid abundance nor the rate of phospholipid synthesis is affected by increased long-chain acyl-thioester concentrations caused by conversion of exogenous fatty acids to acyl-CoA. To stabilize phospholipid abundance, we created an inhibition function *f*([total PL]) that reduces the

LPA synthesis reaction rate according to the abundance of total phospholipids accumulated **(Equation S4)**:

**(Eq. S4)**

$$f([\text{total PL}]) = \frac{1}{1+\frac{[\text{total PL}]}{Ki}}$$

This introduces feedback inhibition of PlsB by phospholipid accumulation. The feedback inhibition effectively stabilizes flux through PlsB despite the stepwise increase in acyl-CoA **(Figure S1)**.

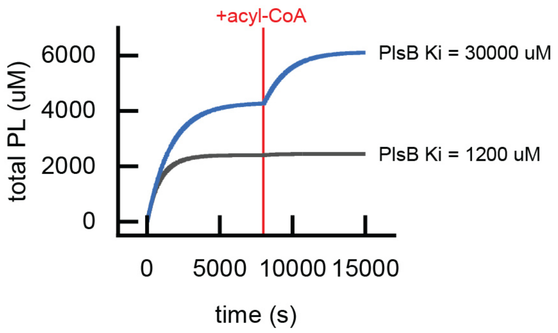

**Figure S1.** Phospholipid abundance from initiation of the simulation to the stepwise increase of acyl-CoA at 8000 s. Implementing feedback inhibition on PlsB activity by total phospholipid content stabilizes phospholipid abundance despite the increase of all substrates triggered by acyl-CoA (depicted by black line generated from setting the PlsB inhibition constant in **Equation S4** to 1200 µM). Removing feedback by increasing the inhibition term $K_I$ to 30000 µM (blue line) allows acyl-CoA synthesis to increase phospholipid abundance.

In the model, PlsB binds and reacts with C16:0, C18:0, C16:1, and C18:1 thioesters; however PlsB does not react efficiently with C16:1 thioesters (3). To implement substrate preferences for PlsB, a coefficient was introduced for the C16:1 thioester binding constant. All PlsB-substrate complexes reacted at identical rates.

#### Substrate competition for PlsC and PA synthesis.

Acyl-thioester substrate competition for PlsC was implemented similarly as for PlsB **(Scheme S2)**. In the model, PlsC binds and reacts with C16:0, C16:1, C18:1, and C20:1 thioesters but does not react efficiently with C16:0 thioesters. Thus a coefficient was introduced for the binding constant for C16:0 thioester species to reduce substrate binding relative to preferred substrates.

$$\text{acyl-thioester + PlsC} \underset{k_{\text{off}}}{\overset{k_{\text{on}}}{\rightleftharpoons}} \text{PlsC-substrate complex + LPA} \xrightarrow{v_{\text{PlsC}}} \text{PlsC + PA (+ holo-ACP)}$$

**(Scheme S2)**

All PlsC-acyl-thioester enzyme-substrate complex subsequently reacted with each individual LPA species according to a standard Michaelis-Menten kinetic function $v_{\text{PlsC}}$ **(Equation S5)**:

**(Eq. S5)** $$v_{\text{PlsC}} = \frac{k_{cat}*[\text{PlsC-substrate complex}]*[\text{LPA}]}{K_M+[\text{LPA}]}$$

#### Phospholipid synthesis and phospholipid dilution by simulated growth.

For simplicity, conversion of each individual PA species to its corresponding membrane phospholipid was simulated as a single step labelled as “CdsA”. This combines PA conversion to CDP-DAG by CdsA and all subsequent pathways for generation of PG and PE. The reaction rate $v_{PL}$ converting each individual PA species to its corresponding phospholipid was simulated as a single-substrate Michaelis-Menten reaction **(Equation S6)**:

**(Eq. S6)** $$v_{\text{PL}} = \frac{k_{cat}*[\text{CdsA}]*[\text{PA}]}{K_M+[\text{PA}]}$$

To record acyl chain distributions in simulated membrane phospholipids and account for total phospholipid abundance (which allows its use as a feedback inhibition signal controlling PlsB), phospholipids produced by the “CdsA” reaction with PA were steadily depleted by the model according to simple mass action kinetics **(Equation S7)**:

**(Eq. S7)** $$v_{\text{PL dilution}} = k_{growth} * [\text{PL}]$$

| **Table S2.** Reaction parameters used. | |
|---|---|
| **ACC-FabD initiation (Equation S1)** | |
| Vmax | 100 μM/sec |
| Kma (for acetyl-CoA) | 500 μM |
| Kmb (for holo-ACP) | 10 μM |
| Ki | 5 μM |
| **FabH (Equation S2)** | |
| kcat | 5/sec |
| [FabH] | 1.6 μM |
| Kma (for acetyl-CoA) | 10 μM |
| Kmb (for malonyl-ACP) | 2 μM |
| **FabB (Equation S3)** | |
| kcat | 10/sec |
| [FabB] | 5 μM |
| Kma (for acyl-ACP) | 2.5 μM |
| KmB (for malonyl-ACP) | 2.5 μM |
| c(C4:0-ACP) | 0.33 |
| c(C6:0-ACP) | 0.86 |
| c(C8:0-ACP) | 0.97 |
| c(C10:0-ACP) to C12:0-ACP (saturated branch) | 0.97 |
| c(C10:0-ACP) to C12:1-ACP (unsaturated branch) | 2.5*c(C10:0-ACP) |
| c(C12:0-ACP) | 1.0 |
| c(C14:0-ACP) | 0.3 |
| c(C16:0-ACP) | 0.02 |
| c(C12:1-ACP) | 1.0 |
| c(C14:1-ACP) | 0.3 |
| c(C16:1-ACP) | 0.15 |
| c(C18:1-ACP) | 0.01 |
| **PlsB substrate binding, catalysis, and inhibition (Scheme S1, Equation S4)** | |
| kon | 1000/(μM*s) |
| koff | 100/s |
| kon(16:1) | 0.05*kon |
| kcat | 10/s |
| Ki | 1200 μM |
| **PlsC substrate binding and catalysis (Scheme S2, Equation S5)** | |
| kon | 1000/(μM*s) |
| koff | 100/s |
| kon(16:0) | 0.05*kon |
| kcat | 10/s |
| Km | 0.15 μM |
| **CdsA (phospholipid synthesis from PA) (Equation S6)** | |
| kcat | 10/s |
| [CdsA] | 0.2 μM |
| Km | 10 μM |
| **Phospholipid dilution by growth (Equation S7)** | |
| kgrowth | 0.0006/s |
